# Volatile-Mediated Plant Defense Networks: Field Evidence for Isoprene as a Short-Distance Immune Signal

**DOI:** 10.1101/2025.02.03.636237

**Authors:** Peiyuan Zhu, Baris Weber, Maaria Rosenkranz, Andrea Polle, Andrea Ghirardo, Jan Muhr, A. Corina Vlot, Jörg-Peter Schnitzler

**Author notes:** Corresponding author: Jörg-Peter Schnitzler, **Email:**. **Author Contributions:** P.Z., M.R., A.P., and J.-P.S. designed research; P.Z. performed research with B.W. (GC-MS) and J.M. (greenhouse); A.C.V. contributed reagents; P.Z. analyzed data; P.Z., M.R., A.P., A.G., A.C.V., and J.-P.S. interpreted data; and P.Z. wrote the paper with input from all authors. **Competing Interest Statement:** The authors declare no competing interests.

## Abstract

Isoprene, the most abundant hydrocarbon emitted by vegetation, protects photosynthesis against oxidative and thermal stress and significantly impacts atmospheric chemistry. While laboratory studies suggest a role for isoprene in plant defense, its function in plant-plant communication under natural conditions remains unclear. Here, we demonstrate that isoprene acts as a signaling molecule, triggering systemic immune responses in neighboring plants against bacterial pathogens under field conditions. We established an experimental system using wild-type and transgenic silver birch (*Betula pendula*) lines engineered to emit varying levels of isoprene. Over two growing seasons, we examined the effects of birch-emitted volatile organic compounds on neighboring Arabidopsis thaliana, including wild-type and immune signaling mutants (*llp1*: legume lectin-like protein 1; *jar1*: jasmonate resistant 1). Isoprene emission rates positively correlated with systemic resistance in receiver plants, with higher emissions enhancing inhibition of *Pseudomonas syringae* growth. This immune response occurred independently of jasmonate signaling but required functional LLP1, suggesting a specific recognition pathway. By combining *Arabidopsis* mutant responses with birch volatile profiles, we confirmed that isoprene, rather than other terpenoids, mediates this effect. These findings reveal an unrecognized ecological function of isoprene as an airborne immune signal, establishing transgenic birch as a robust model for studying volatile-mediated plant defense networks in natural environments. Our results provide new insights into the molecular mechanisms underlying plant volatile perception and expand our understanding of chemical communication in terrestrial ecosystems.

**Significance Statement:** Isoprene, the most abundant volatile organic compound released by plants, protects against abiotic stress. Here, we demonstrate that isoprene also functions under ambient atmosphere as a short-distance immune signal, enhancing neighboring plants’ resistance to bacterial pathogens. This finding reveals a putative ecological role of isoprene in plant defense networks and suggests strategies for sustainable agriculture and forest management under environmental stress.

## Introduction

Plants are involved in complex interactions with their environment, employing sophisticated mechanisms to respond to stress and communicate with other organisms. The production and emission of biogenic volatile organic compounds (BVOCs) play a central role in plant ecology and defense (1, 2). BVOCs include a wide range of volatile organic molecules released by plants, such as isoprene, monoterpenes, sesquiterpenes, and other compounds (3, 4). These volatiles attract pollinators and seed dispersers, repel herbivores and pathogens, and facilitate plant-to-plant communication (4–7).

Terpenoids constitute over 50% of BVOCs (8). This group includes isoprene (C_5_), monoterpenes (C_10_), and sesquiterpenes (C_15_). Isoprene is the most abundant BVOC, with an estimated emission of 500 Tg carbon per year, accounting for nearly half of the total carbon released as BVOCs. Monoterpenes and sesquiterpenes contribute 15% and 3%, respectively(9, 10). Various terpenoids compounds have been shown to play roles in plant-environment interactions (11, 12). For example, different monoterpenes and sesquiterpenes engage in various direct and indirect plant defenses against herbivores and pathogens, as well as in plant-plant communication (13, 5, 14). Recent research has shown that monoterpenes and sesquiterpenes released by sender plants can induce immunity in receiver plants (15–17). Beyond plant defense, monoterpenes and sesquiterpenes significantly influence ecosystem-level processes and play crucial roles in atmospheric chemistry through their interactions with other compounds and effects on air quality (18, 19).

Despite extensive research on terpenoids in plant defense, the function of isoprene, the smallest and most abundant terpenoid, remains debated. Studies have shown that isoprene protects plants from abiotic stress, particularly high temperature and oxidative stress (20–22). Its thermo-protective effects are attributed to thylakoid membrane stabilization and modification of reactive oxygen species (ROS) (23, 24). Recent studies on plants with modified isoprene emission capacity suggest that isoprene indirectly affects stress protection by altering gene expression, protein abundance, and internal signaling pathways (21, 25, 26). Externally perceived isoprene has also been shown to induce receiver plant immunity by altering plant defense pathways (16) and upregulating stress-responsive genes and transcription factors in receiver plants (27). Terpenoids’ function in inducing plant immunity is linked to the salicylic acid (SA) and jasmonic acid (JA) signaling pathways (28, 29). SA is typically associated with defense against biotrophic pathogens, while JA is associated with defense against necrotrophic pathogens and herbivores (30–32). Studies on monoterpenes and sesquiterpenes show that different compounds interact with specific phytohormone pathways to enhance plant resistance to pathogens and herbivores. For example, β-caryophyllene, a sesquiterpene, induces resistance to bacterial pathogens in *Arabidopsis thaliana* by modulating JA pathways (16), whereas monoterpenes such as α-pinene and β-pinene, as well as isoprene, can induce defense mechanisms dependent on SA biosynthesis and signaling (17, 16). Recent data suggest that monoterpenes can act as signaling molecules in plant-plant communication(33). While specific monoterpenes (α-pinene, β-pinene) and sesquiterpenes (β-caryophyllene) roles in plant defense signaling are well-characterized, the molecular mechanisms underlying isoprene-mediated plant immunity and its potential interactions with other defense-related terpenoids remain to be elucidated.

Translating laboratory study results on BVOC emissions to field conditions is a major challenge and a significant gap in understanding plant-environment interactions under natural conditions. Natural environments’ complexity, including variable temperatures, light intensities, and biological interactions, can strongly affect BVOC emission rates and plant responses (2, 34). Despite recent advances in understanding isoprene’s function in plant defense against biotic stress, few studies link findings from controlled laboratory experiments to natural conditions (16, 35, 36). Transgenic silver birch (*Betula pendula* Roth) (37) represents a unique model system to address these challenges. Silver birch, an ecologically and economically important tree species, emits mainly monoterpenes and sesquiterpenes and little isoprene (38, 39). Silver birch plays a key role in boreal and temperate forest ecosystems, influencing nutrient cycling, biodiversity, and atmospheric chemistry (40, 41). Transgenic silver birch lines with modified isoprene emission profiles provide a unique opportunity to investigate the effects of altered BVOC emissions on plant-environment interactions and neighboring plants in a natural environment (42, 37).

The aim of this study was to determine whether plant-released isoprene affects the immunity of neighboring plants under natural conditions. We used transgenic birch lines that emit different amounts of isoprene (37). We employed *Arabidopsis thaliana* as receiver plants and tested their immune response by their ability to suppress the growth of pathogenic bacteria (*Pseudomonas syringae*). In addition to *Arabidopsis thaliana* wild type, we included isogenic knock-out lines of legume lectin-like protein 1 (*llp1*) and jasmonate resistance 1 (*jar1*) because these gene functions are required to mediate monoterpene-(LLP1) and sesquiterpene-induced (JAR1) immunity, respectively (43, 33, 16). Hence, the *Arabidopsis* mutants are excellent control plants to test the efficacy of specific signaling cues. Using transgenic birch trees, and different *Arabidopsis* mutants in an outdoor setting over two consecutive growing seasons, this study evaluates the role of isoprene in enhancing plant immunity in a natural environment and provides a better understanding of plant-to-plant communication in natural ecosystems.

We hypothesize that isoprene emitted by silver birch in a natural environment enhances the immunity of neighboring plants against pathogens. This ecological interaction could represent a novel mechanism of plant-to-plant communication in forest ecosystems. Using Arabidopsis wild-type and defense signaling mutants (*llp1* and *jar1*), this study investigates the role of isoprene as an airborne immune signal under natural conditions.

## Results

### Terpenoid Emission Profiles of Transgenic Silver Birch Lines

Transgenic modifications profoundly altered terpenoid emissions across different compound classes, with patterns persisting over two growing seasons (Fig. 2). Isoprene emissions showed clear genotypic effects. Our 2022 data demonstrated that birch line 12 emitted the highest levels, with line 03 showing intermediate levels (P < 0.05), and WT maintaining the lowest emissions (Fig. 2a). We observed the same trend in 2023, where both line 03 and line 12 significantly higher than WT, while line 06 showed intermediate levels (Fig. 2b; *H₂* = 10.231, *df* = 3, *P* < 0.05, *η²* = 0.425).

**Figure 1.**
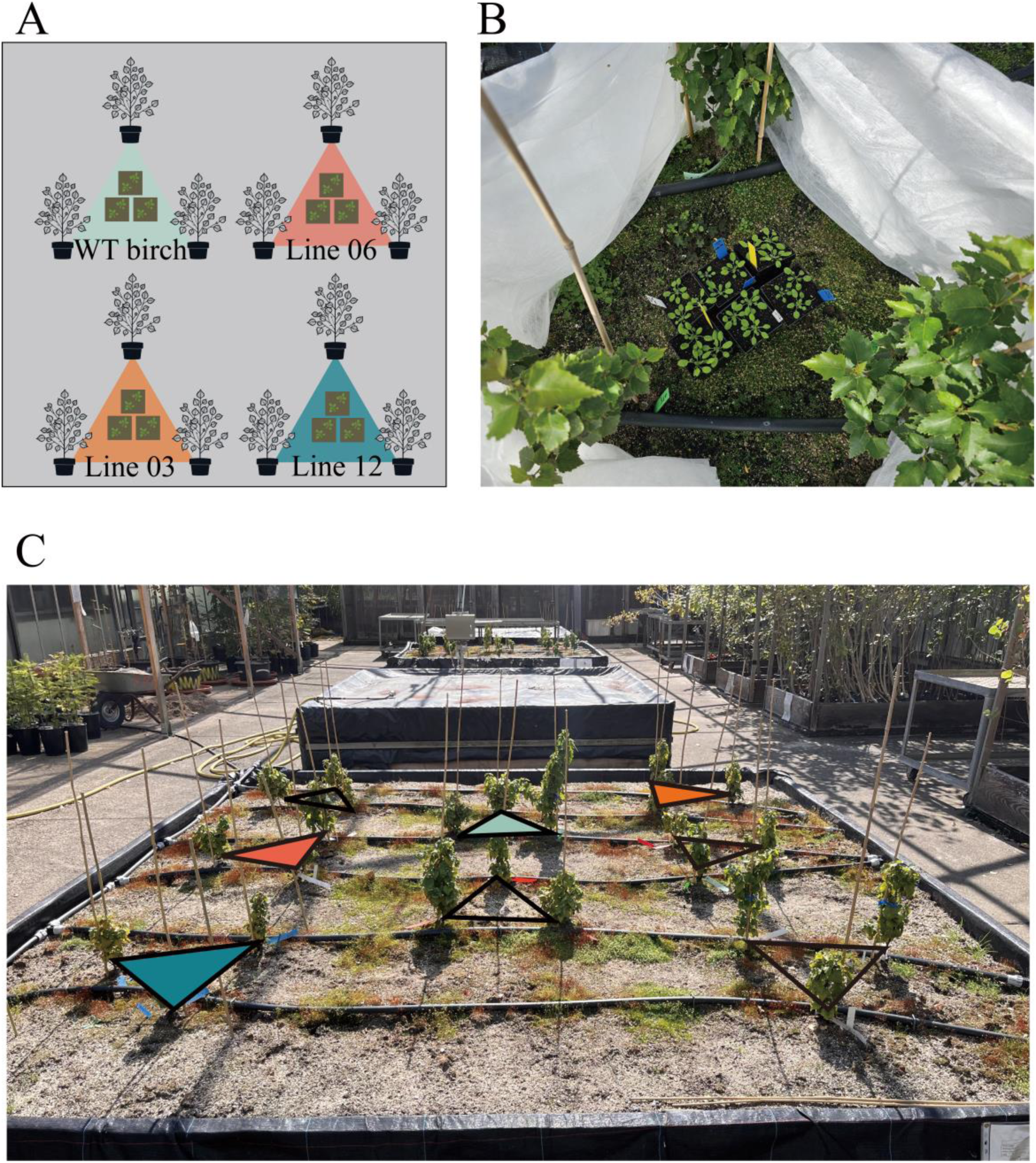
Experimental design for studying isoprene emission effects from transgenic silver birch on Arabidopsis thaliana. (A) Schematic representation of silver birch (*Betula pendula*) planting arrangement showing wild-type (WT) and transgenic lines (06, 03, and 12) in triangular formations. (B) Magnified view of the central experimental area within a birch triangle, showing Arabidopsis thaliana plants (Col-0 wild-type, *llp1*, and *jar1* mutants) positioned for VOC exposure. (C) Field implementation of the experimental setup in an outdoor open-air greenhouse showing the spatial arrangement of birch triangles with colored markers indicating different transgenic lines.

**Figure 2.**
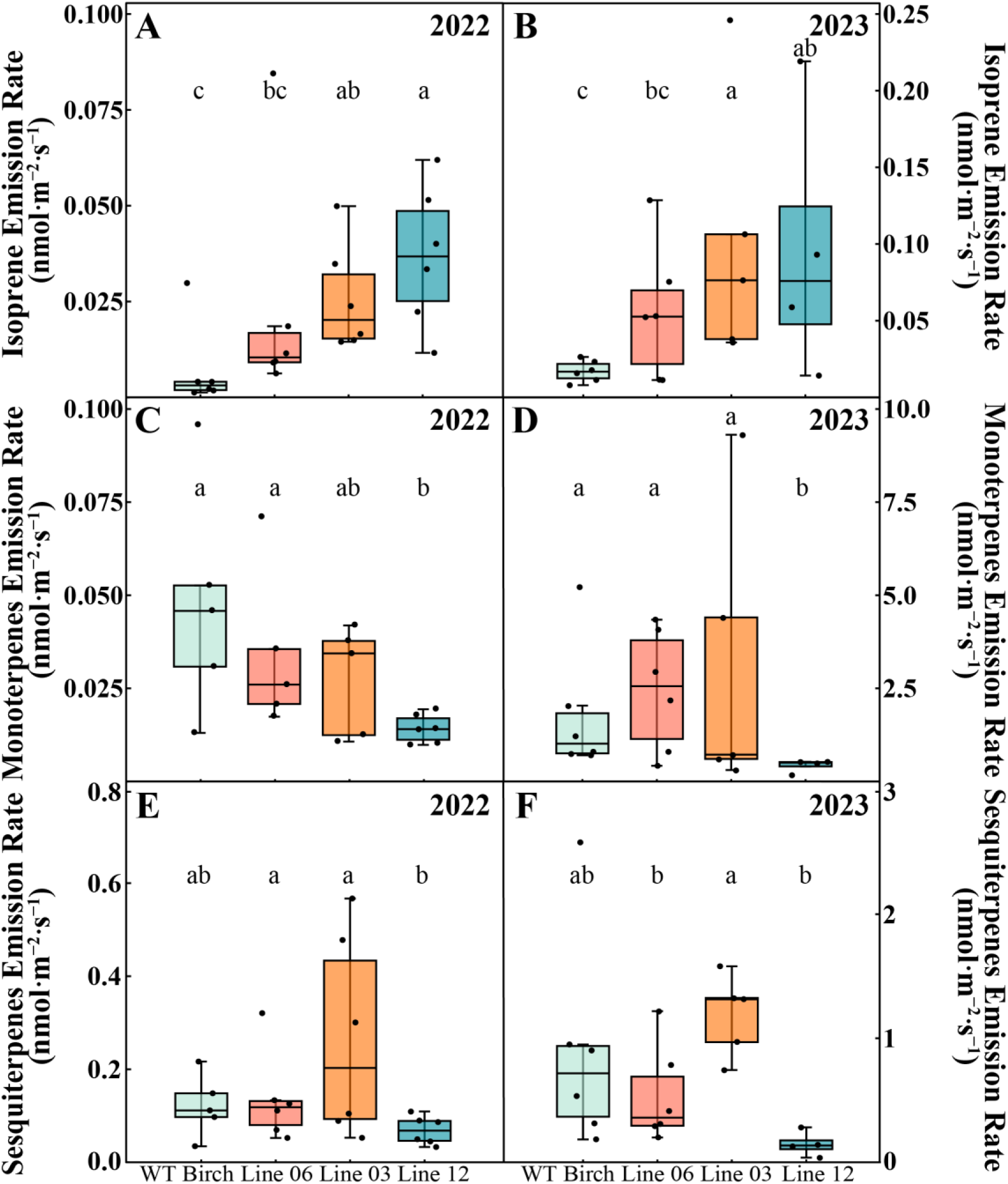
Volatile terpenoid emission profiles from wild-type and transgenic silver birch lines. Emission rates of (A, B) isoprene, (C, D) monoterpenes, and (E, F) sesquiterpenes measured from wild-type (WT) *Betula pendula* and transgenic lines (06, 03, and 12) during growing seasons of 2022 (A, C, E) and 2023 (B, D, F). Box plots show median (horizontal line), interquartile range (box: 25th-75th percentiles), and 10th-90th percentiles (whiskers); *N* ≥ 4 biological replicates. Different letters indicate significant differences between lines (*P* < 0.05, Kruskal-Wallis test with Dunn’s post-hoc comparison).

Monoterpenes showed a reverse emission pattern compared to isoprene. In 2022, WT birch and line 06 were our highest emitters (Fig. 2c), while line 12 showed markedly reduced emissions (*P* < 0.05). When we repeated the measurements in 2023, we detected higher overall emissions, but line 12 still maintained its low-emission characteristic (Fig. 2d; *H₂* = 7.198, *df* = 3, *P* < 0.1).

In 2023, emissions of sesquiterpenoids exhibited significant genotype-specific differences (*H_2_* = 8.703, *df* = 3, *P* < 0.05, *η^2^* = 0.34), with notably higher emissions observed in line 03 compared to lines 06 and 12 (0.517 ± 0.152 and 0.236 ± 0.105 nmol m^-2^·s^-1^, respectively) (1.157 ± 0.146 nmol m^-²^ s^-1^).

While the relative emission patterns between the birch genotypes stayed consistent across the years, absolute emission rates were higher in 2023, corresponding with higher sampling temperatures (Supplementary Fig. S4).

### Isoprene from *Betula pendula* Affects *Arabidopsis* Immune Responses

GC-MS analysis revealed that transgenic silver birch lines exhibited distinct isoprene emission patterns, with lines 03 and 12 showing consistently higher emissions compared to WT and line 06 across both 2022 and 2023 (Figs. 2a and 2b). When these high isoprene-emitting lines were tested for their ability to induce plant immunity and inhibit bacterial growth on WT *Arabidopsis*, we observed a strong positive correlation (*R²* = 0.7881, *P* = 0.0032). Across both years, exposure to volatiles from transgenic birch lines showed varying degrees of bacterial growth inhibition in WT *Arabidopsis*, with line 03 consistently showing significant effects (*P* < 0.05), while line 12’s effect was stronger in 2022 compared to 2023 (Fig. 3a and 3b). Line 06, which exhibited intermediate isoprene emission levels, showed moderate bacterial growth inhibition that was statistically intermediate between WT birch and high-emitting lines, consistent with its intermediate isoprene emission profile.

**Figure 3.**
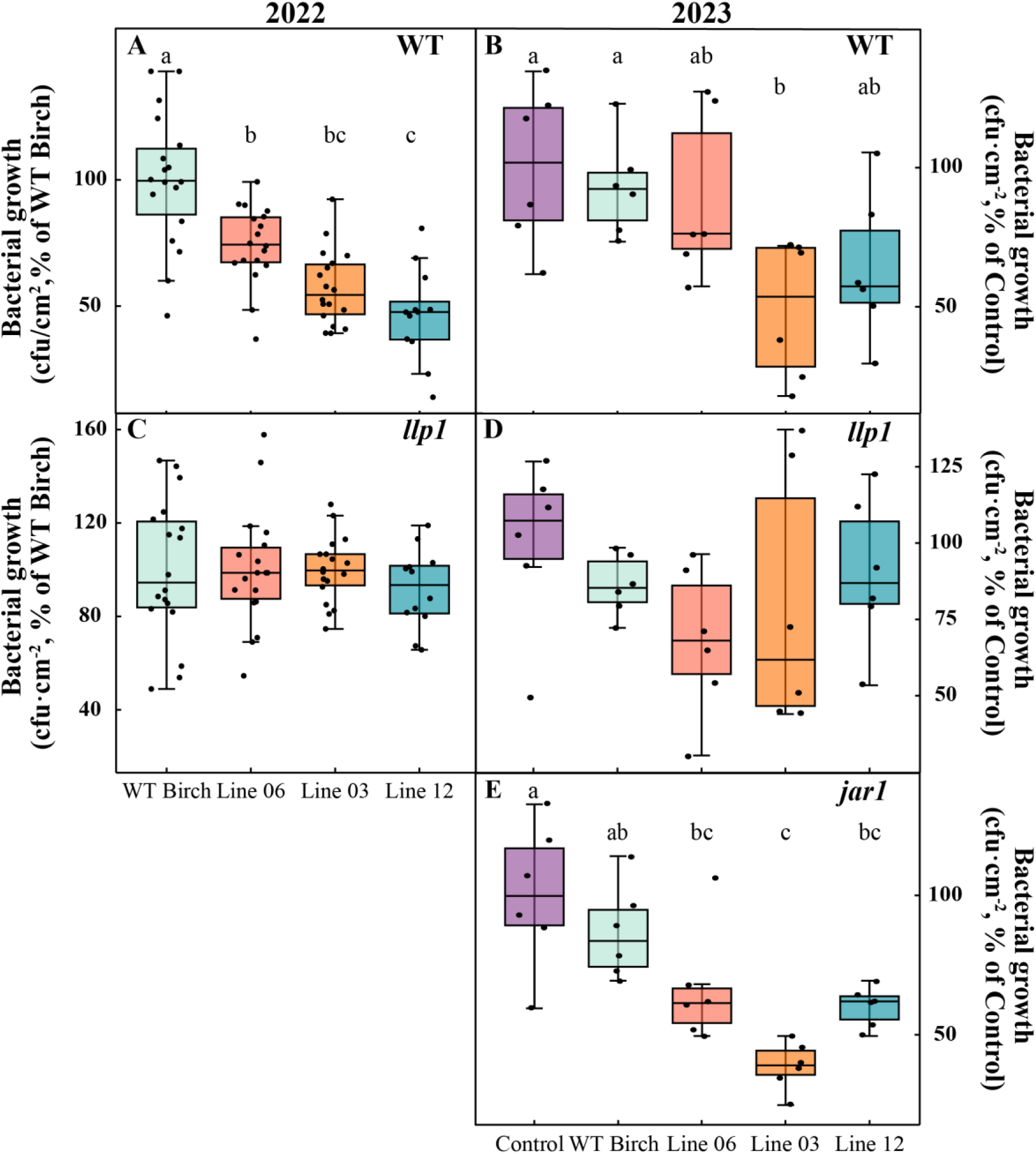
Impact of silver birch volatile emissions on bacterial growth in wild-type and mutant *Arabidopsis thaliana* (*llp1* and *jar1*). Bacterial growth quantification in (A, B) wild-type (WT), (C, D) *llp1* mutant, and (E) *jar1* mutant *A. thaliana* plants exposed to VOCs from wild-type and transgenic *Betula pendula* lines (06, 03, and 12) during 2022 (A, C) and 2023 (B, D, E). Bacterial growth is expressed as percentage relative to plants exposed to WT birch emissions. Box plots show median (horizontal line), interquartile range (box: 25th-75th percentiles), and 10th-90th percentiles (whiskers); individual data points represent independent biological replicates (*N* ≥ 6). Different letters indicate significant differences between treatments (*P* < 0.05, one-way ANOVA).

In 2023, we included ambient air as an additional control, WT *Arabidopsis* exposed to ambient air showed high bacterial growth levels comparable to some treatments, though significantly higher than those exposed to high isoprene-emitting lines. Despite birch line 12 showing consistently lower monoterpenoid and sesquiterpenoid emissions compared to other genotypes (Fig. 2c-f), its elevated isoprene emissions corresponded with strong bacterial growth inhibition in receiver plants, suggesting isoprene’s specific role in this response.

To quantitatively analyze the relationship between isoprene emission and bacterial growth inhibition, we performed correlation analysis using the actual isoprene emission rates and bacterial suppression levels measured for each experimental group in 2023 (Fig. 4a). Due to varying emission patterns among individual birch trees (Fig. 2 and Supplementary Fig. 1), the average isoprene emission rate was calculated for each experimental group surrounding the *Arabidopsis* receivers. Statistical analyses revealed significant differences (*P* < 0.05) among the experimental triangles and a substantial positive correlation between isoprene emission levels and bacterial growth inhibition in the receiver plants. Notably, the correlation analyses demonstrated no significant relationship between monoterpene or sesquiterpene emissions and bacterial growth inhibition on any genotype (Supplementary Fig. 3), thereby identifying isoprene as the primary volatile signal inducing plant immunity in neighboring receivers.

**Figure 4.**
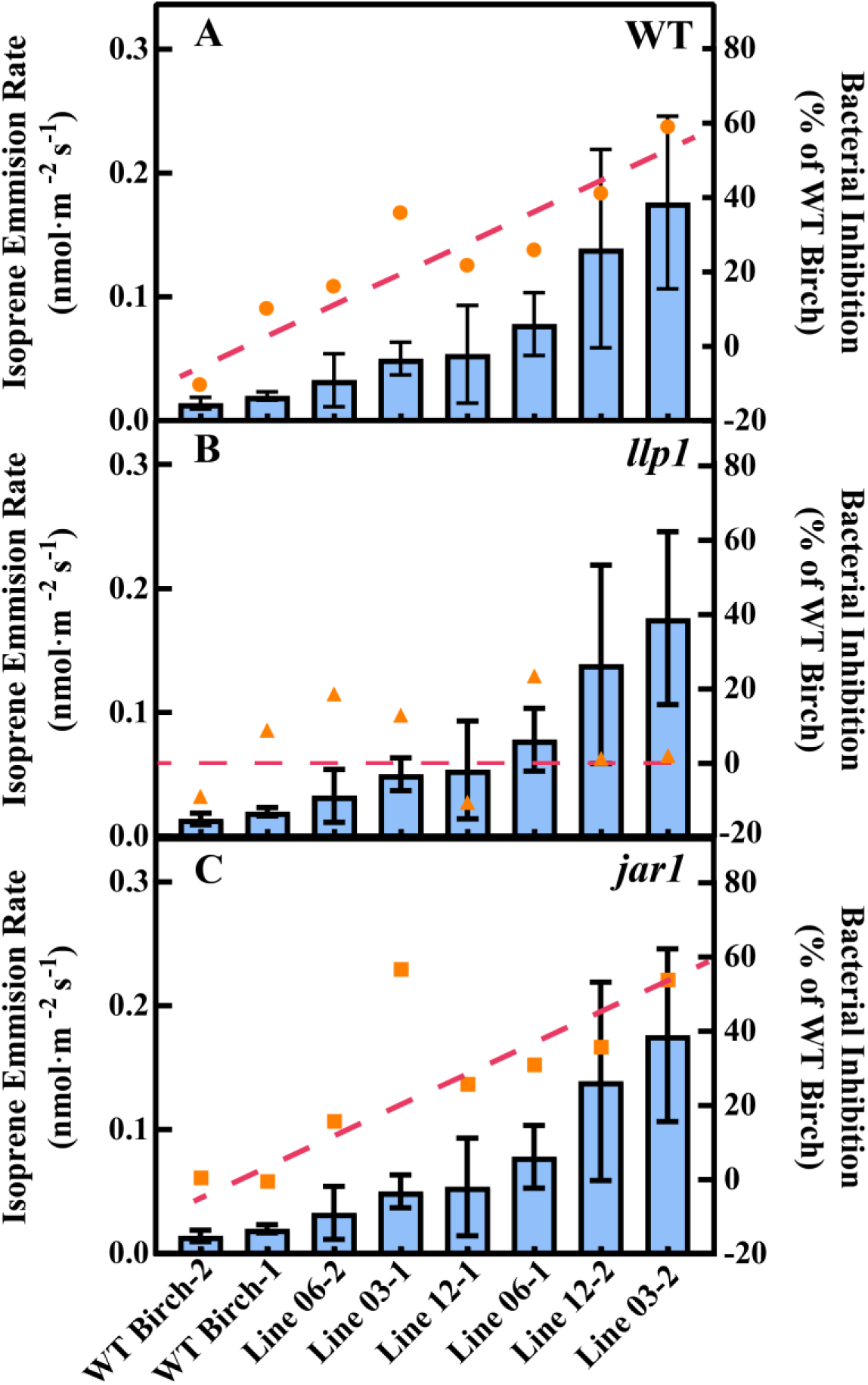
Correlation between isoprene emission rates and bacterial inhibition for each triangle in wild-type (WT) and mutant birch lines with defective defense signaling pathways. Isoprene emission rates (blue bars, mean ± SEM) and bacterial inhibition (orange markers, % of WT Birch) are presented for (A) wild-type (WT), (B) *llp1* mutant (defective in monoterpene-mediated salicylic acid defense signaling), and (C) *jar1* mutant (impaired in sesquiterpene-mediated jasmonic acid defense signaling) birch lines. Red dashed lines indicate the linear regression between emission rates and bacterial inhibition. Statistical analysis revealed a significant positive correlation in WT *Arabidopsis* (*R²* = 0.7881, *P* = 0.0032). In contrast, no significant correlation was observed for *llp1* (*R²* = 0.0094, *P* = 0.8195) or *jar1* (*R²* = 0.4908, *P* = 0.0529) mutant lines. Numbers following line designations represent different emission triangles.

### Impact of Birch VOCs on Bacterial Growth in *Arabidopsis* Mutants

To further understand the effects of silver birch VOCs on plant immunity in *Arabidopsis* receivers, we examined bacterial growth responses in two *Arabidopsis* mutants with defects in distinct defense signaling pathways: *jar1* (impaired in jasmonate signaling) and *llp1* (deficient in monoterpene signaling).

In bacterial growth assays with *llp1* mutants, no significant differences were observed between birch genotypes in either 2022 or 2023 (Fig. 3c and 3d). Correlation analyses revealed no significant relationships between any terpenoid emissions and bacterial growth inhibition in these plants Correlation analyses revealed no significant relationships between any terpenoid emissions and bacterial growth inhibition in these plants (P > 0.05; Fig. 4b and Supplementary Fig. 3). LLP1, which functions as a component of systemic acquired resistance, appears to be required for the plant’s response to volatile signals from birch. In contrast, *jar1* mutants displayed genotype-specific responses to birch VOCs in 2023 (Fig. 3e). High isoprene-emitting lines 03 and 12 induced significant bacterial growth suppression in *jar1* plants compared to ambient air controls and WT birch exposure (*P* < 0.05). Correlation analysis revealed a moderate positive relationship between isoprene emission rates and bacterial growth inhibition in *jar1* mutants (*R²* = 0.4908, *P* = 0.0529; Fig. 4c), while neither monoterpenes (*R²* = 0.0500, *P* = 0.5947) nor sesquiterpenes (R² = 0.0349, P = 0.6577) showed significant correlations (Supplementary Fig 3e and 3f). The preserved response in *jar1* mutants demonstrates that isoprene-induced immunity operates independently of the jasmonate signaling pathway. While the specific molecular mechanisms remain to be fully elucidated, these results support the role of isoprene as a distinct volatile signal capable of triggering plant immunity through pathways that are separate from classical jasmonate-mediated defense signaling.

Together, these mutant analyses suggest distinct signaling requirements for different VOCs in triggering plant immunity, highlighting the complexity of volatile-mediated defense responses in plants.

## Discussion

Our experimental design, using transgenic silver birch trees with modified terpenoid emission profiles and *Arabidopsis* plants as receivers, demonstrates the efficacy of isoprene-mediated plant immunity under natural conditions. The consistent response patterns observed over two years under different environmental conditions underline the robustness of this plant-to-plant communication system. While higher temperatures in 2023 were associated with higher isoprene emissions (see Supplementary Figures 2 and 4), the basic relationships between VOC emissions and bacterial resistance remained stable, indicating the reliability of this defense mechanism under different environmental conditions. Recent studies have further demonstrated the critical role of temperature and light intensity in modulating VOC emissions and plant responses under field conditions (44, 45). This is particularly true for isoprene, whose biosynthesis and emission respond rapidly to current temperature and light conditions, with emission rates closely tracking environmental changes over short time periods (46, 47).

The constant VOC emission patterns of the birch mutants over two years led us to investigate the metabolic relationships within terpenoid biosynthesis. We observed an inverse relationship between isoprene and monoterpene emission rates, suggesting a metabolic trade-off in the plastidic methyl-D-erythritol phosphate (MEP) pathway. This crosstalk in emission rates likely reflects multiple regulatory mechanisms, including competition for common precursors (IPP and DMAPP), differential enzyme kinetics, and transcriptional regulation of the respective biosynthetic genes in the MEP pathway (48, 49). The complex interactions between introduced genes and the plant’s native genome can lead to unexpected phenotypes or metabolic shifts (50, 51). While increased isoprene emission is often correlated with decreased emission of monoterpenes and sesquiterpenes, birch line 03 showed high emission of both terpenoid groups, indicating complex metabolic adaptations that warrant further investigation (52, 53).

Our results may have important ecological implications beyond the responses of individual plants. The increase in microbial pathogen resistance in neighboring *Arabidopsis* plants suggests that volatile signals play an important role in plant-plant communication. This communication may be particularly relevant in densely vegetated, low-stature ecosystems like the Arctic and Sub-Arctic tundra, where plants grow in close proximity (54, 55). Similarly, in tropical ecosystems, such as hyper dominant species in the Amazon (*Inga*, *Protium*), demonstrate isoprene’s role in enhancing resilience to environmental stress (56, 57). In these ecosystems, isoprene emission is linked to thermotolerance and oxidative stress mitigation, facilitating coexistence and communication in diverse communities. These findings extend our understanding of the ecological implications of terpenoid emissions, suggesting that isoprene-mediated signaling is not only critical for individual plant adaptation but also shapes broader community dynamics across ecosystems.

The pathway-specific nature of this communication system is confirmed by the differential responses in the mutant *Arabidopsis* lines. The differential responses observed in WT and mutant *Arabidopsis* lines (*llp1* and *jar1*) suggest that the effects of isoprene on plant immunity involve LLP1, which functions as a parallel signaling component in systemic acquired resistance (SAR) (33, 16). Similarly, monoterpene-induced immunity relies on functional LLP1, suggesting overlapping responses to isoprene and monoterpenes via the SAR regulatory network (17, 33, 16). In contrast, the preserved responsiveness of *jar1* mutants suggests that JA-dependent defenses may play a minor role in this process. While isoprene emissions correlated significantly with bacterial growth inhibition (Fig. 4), neither monoterpene nor sesquiterpene emissions showed similar correlations in our experimental setup (Supplementary Fig. 3). These results highlight the complexity of plant immune responses to VOCs and suggest that plants may use multiple pathways to enhance immunity (58). The mechanism by which plants perceive isoprene as a signal is still mostly unclear. The results presented here suggest that the receiver plant processes a perceived isoprene signal via the SAR signaling cascade. Additionally, laboratory experiments using *Arabidopsis* overexpressing the isoprene synthase gene from *Eucalyptus* have shown that signaling networks associated with specific phytohormones such as gibberellic acid and stress tolerance are altered by isoprene production (59). This highlights the need for future studies, ideally under natural conditions as here, to include additional defense-related mutants beyond *llp1* and *jar1*, particularly those involved in MAPK cascades and calcium signaling, to gain deeper insight into the integration of isoprene into broader immune networks (60–62). Furthermore, multi-omics approaches such as RNA-Seq, untargeted metabolomics, and post-translational modification (PTM) proteomics could elucidate the molecular mechanisms underlying VOC-mediated interactions and identify key regulatory hubs (63–65). Recent studies on ectomycorrhizal fungal-plant interactions demonstrate the potential of these integrative approaches to dissect plant VOC signaling networks (66). Our study demonstrates that isoprene can enhance plant immunity over short distances (<50 cm) under natural conditions, adding to its known benefits in protecting photosystems against thermal stress (67, 68) and oxidative damage (69, 23). However, these beneficial plant-level effects must be weighed against isoprene’s atmospheric chemistry impacts, particularly its contribution to ozone formation in NOx-rich environments (70) and secondary organic aerosol formation (64, 71). Based on these findings, we propose a context-dependent approach to managing isoprene-emitting species: In agricultural settings and natural ecosystems distant from urban centers, the benefits of isoprene emission for plant health and community resilience may outweigh atmospheric concerns. However, in urban and suburban areas where NOx pollution is prevalent, priority should be given to low-isoprene-emitting species to minimize air quality impacts. This spatial optimization strategy allows us to harness the beneficial effects of isoprene for plant health while minimizing its negative atmospheric impacts in pollution-sensitive areas.

The present evidence for isoprene as a VOC that stimulates the plant immune system is one more step towards answering the question raised by Sharkey and Singsaas (1995): Why do plants emit isoprene? Future research on isoprene as a signal molecule should focus on two key aspects. First, we need to understand the quantitative relationships between isoprene concentration and immune activation in recipient plants under natural systems. Second, it is crucial to determine how environmental factors like temperature, light intensity, and humidity modulate this defense mechanism. Long-term studies that consider soil properties, microbiome dynamics, and atmospheric chemistry will be crucial for predicting ecosystem responses to isoprene.

## Materials and Methods

### Plant and Microbial Materials and Growth Conditions

In the experiment, three genetically modified isoprene-overexpressing lines of Finnish elite birch (lines 03, 06 and 12) clones were used alongside wild-type Finnish elite birch (*Betula pendula* Roth). The transformation and cultivation procedures of these transgenic plants have been described previously (37). In autumn 2021, the 4-year-old birch plants were transferred from the phytochambers to an outdoor, caged area of the Department of Forest Botany and Tree Physiology at the University of Göttingen, Germany. The young trees were grown in triangles of the same genotype at 50 cm distance in large soil-filled boxes (Fig. 1a-c). The plants over-wintered outdoors and were used in subsequent years for the experiments.

In addition, wild-type *Arabidopsis thaliana* (ecotype Columbia-0 (Col-0)) and the isogenic *Arabidopsis* KO mutant lines legume lectin-like protein 1 (*llp1*) and jasmonate resistant 1 (*jar1*) were used. *Arabidopsis* plants were grown in climate-controlled chambers in a soil mixture containing a 1:5 ratio of coarse sand to normal soil. Growth conditions were maintained at a temperature of 20/16°C and a relative humidity of 65/80% (day/night); light irradiation was provided at 100 μmol photons m^-2^ s^-1^ photosynthetic photon flux density (PPFD) with a 10-hour photoperiod. All *Arabidopsis* plants used in the experiments were 4.5 weeks old. *Arabidopsis* plants were not pre-conditioned to outdoor conditions before the experiments.

The bacterial strain used in the infection assays was *Pseudomonas syringae* pv. *tomato* (Pst) DC3000. Bacteria were grown on NYGA medium (0.5% bactoproteose peptone, 0.3% yeast extract, 2% (v:v) glycerol, 1.8% agar, pH 7.0) supplemented with 50 μg·mL^-1^ rifampicin and 50 μg·mL^-1^ kanamycin at 28°C. For inoculation, freshly grown bacteria were suspended in 10 mM MgCl₂ and adjusted to OD₆₀₀ = 0.0002 (approximately 10⁵ colony forming units (CFU) mL^-1^). Two upper leaves of *Arabidopsis* plants were infiltrated with this bacterial suspension using a needleless syringe 84 h after the start of the volatile treatment. Three days after inoculation, *in planta* bacterial titers were determined as follows: three leaf discs per sample were shaken in 10 mM MgCl₂ with 0.01% (v:v) Silwet, serially diluted, plated on NYGA medium and incubated for 2 d at 28°C before colony counting and calculation of *in planta* bacterial titers (33).

### Exposure of *Arabidopsis* to Birch VOCs

To study the effect of isoprene emission on neighboring *Arabidopsis* receiver plants, the experiment was repeated three times independently as biological replicates. For each birch line, two triangular arrangements were established, with six pots of *Arabidopsis* plants (two pots per line: wild type, *llp1*, and *jar1*; three plants per pot) randomly placed in the center area of each triangle (Fig 1. A and C). During sampling, leaf material from the three *Arabidopsis* plants within a pot was pooled to form a single sample, consistent with methods described by Frank *et al*. (2021). To minimize wind disturbance, each triangular setup was enclosed with sticks and non-woven fabric during the 84-hour exposure period (Fig 1. B).

### VOC Collection

For the collection of VOCs, the upper part of each birch tree was covered with a transparent disposable plastic coffee cup (0.5 L), which was then sealed with Teflon bags and tape to ensure an airtight seal. Volatile sampling was performed using two MTS-32 autosamplers (Markes International Ltd, Bridgend, UK) equipped with 2-meter long 0.3 mm PFA tubes attached to each sampling port to enable air sample collection from different locations. The samples were collected by GC glass tubes (Gerstel, Mülheim an der Ruhr, Germany) filled with 60 mg Tenax TA 60/80 and 60 mg Carbopack X 40/60 (both from Sigma-Aldrich, St. Louis, USA). The sample flow rate was 250 ml min^-1^, maintained for 30 min. Prior to sampling, each tube was spiked with 2 µL of δ-2-carene in methanol (429.7 pmol µL^-1^) as an internal standard. After sampling, the GC tubes were stored hermetically sealed at 4°C in a refrigerator for subsequent GC-MS analysis.

### VOC Analyses

The samples were analyzed as previously (54) using thermo-desorption gas chromatography-mass spectrometry (TD-GC-MS; TD by Gerstel; GC type: 7890A, MS type: 5975C, both from Agilent Technologies, Palo Alto, CA, USA) and the following modified GC temperature ramp: initial temperature of 40°C increased at 10°C min^-1^ to 130°C, hold for 5 min, increased to 175°C at 80°C min-1, then to 200°C at 2°C min^-1^, to 220°C at 4°C min^-1^, and finally to 300°C at 100°C min^-1^ and hold for 5 min. Quantitative and qualitative VOC analyses were performed using Agilent GC-MS Enhanced ChemStation, version E.02.00.493 (Agilent, St. Clara, USA) as previously (54), and compounds were annotated by spectral matching using the NIST20 and Wiley (v8) mass spectra libraries and by comparing the calculated non-isothermal Kovats retention indices with those found in the literature (NIST Chemistry, WebBook SRD 69, webbook.nist.gov). Only compounds with a match quality score (ChemStation) greater than 75 were selected for further analysis. Compounds were quantified using calibration curves based on pure standards, and emission rates were calculated based on leaf area and enclosure time (54, 67).

### Statistical Analysis

Orthogonal Partial Least Squares Discriminant Analysis (OPLS-DA) was used to characterize the terpenoid profiles of the different silver birch lines. The pre-processing of the raw GC-MS data consisted of log10 transformation, mean centering and Pareto scaling. Prior to OPLS-DA, principal component analysis (PCA) was performed to identify potential outliers. Samples that fell outside the 95% confidence Hotelling’s T2 ellipse of the PCA were treated as outliers and excluded from further analysis. Subsequently, the volatile emission rates in the data matrix (21 × 38) were set as independent variables (X), while the categorical variables describing the different silver birch lines were set as dependent variables (Y) in the OPLS-DA model. The aim of this analysis was to identify the main terpenoids characteristic of the different silver birch lines. The predictive performance of the model was assessed by the regression sum of squares (R2Y), the prediction sum of squares (Q2Y), the root mean square error of estimation (RMSEE) and the RMSEcv (root mean square error of cross-validation). Models were tested for significance (*P* < 0.05) using the cross-validated ANOVA procedure (68, 69). OPLS-DA score plots were plotted to illustrate the differentiation of birch lines based on their terpenoid emission profiles. All multivariate analyses were performed with the software package SIMCA-P v.13.0.3.0 (Umetrics, Umea, Sweden).

Cell forming unit (CFU) data were first assessed for normality using the Shapiro-Wilk test (*P* > 0.05). One-way analysis of variance (ANOVA) was then conducted to evaluate differences among experimental groups, followed by Tukey’s honestly significant difference (HSD) test for post-hoc pairwise comparisons. For datasets with unequal group sizes, the Tukey-Kramer method was applied. For terpenoid emission rates that did not meet the assumption of normality, the Kruskal-Wallis test was used, followed by Dunn’s test for post-hoc comparisons.

## Acknowledgments

We thank Marvin Blaue from LARI for assistance with climate chamber settings and experimental scheduling, Georg Gerl from EUS for logistical support in transporting research materials between München and Göttingen, and Merle Fastenrath and Cathrin Leibecke (Forest Botany and Tree Physiology, University of Göttingen) for technical assistance.

---

**Fig. S1.**
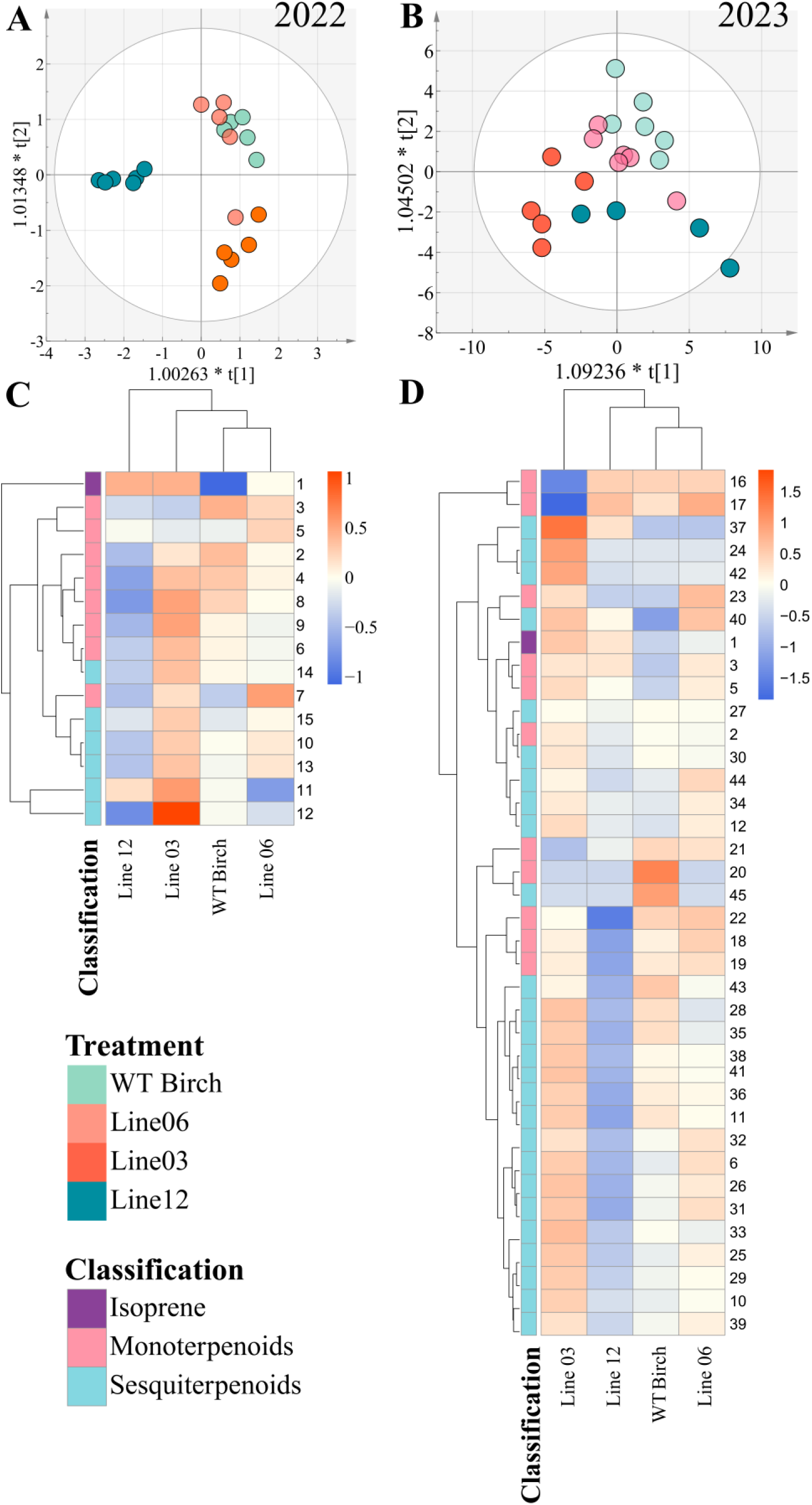
Multivariate analysis of terpenoid emissions from wild-type and transgenic silver birch lines in 2022 and 2023. (A-B) Orthogonal partial least squares discriminant analysis (OPLS-DA) score plots of terpenoid emissions from wild-type (WT) birch and three transgenic lines (06, 03, and 12) for the experiments performed in 2022 (A) and 2023 (B). Each point represents an individual sample. OPLS-DA model fitness: (A) *R^2^X (cum)* = 0.875, *R^2^Y(cum)* = 0.86, *Q^2^Y(cum)* = 0.752, using predictive and two orthogonal components. CV-ANOVA, *P* = 0.0002. (B)*R^2^X (cum)* = 0.752, *R^2^Y(cum)* = 0.898, *Q^2^Y(cum)* = 0.626, using three predictive and two orthogonal components. CV-ANOVA, *P* = 0.003. (C-D) Clustered heatmaps showing the relative abundance (means of *N* ≥ 4) of individual terpenoids (rows) across different birch lines (columns) for 2022 (C) and 2023 (D). Dendrograms on the left show the hierarchical clustering of terpenoids based on their emission patterns, using Euclidean distance and complete linkage. Numbers on the right of heatmap (C, D) correspond to individual terpenoid compounds (Table S1).

**Fig. S2.**
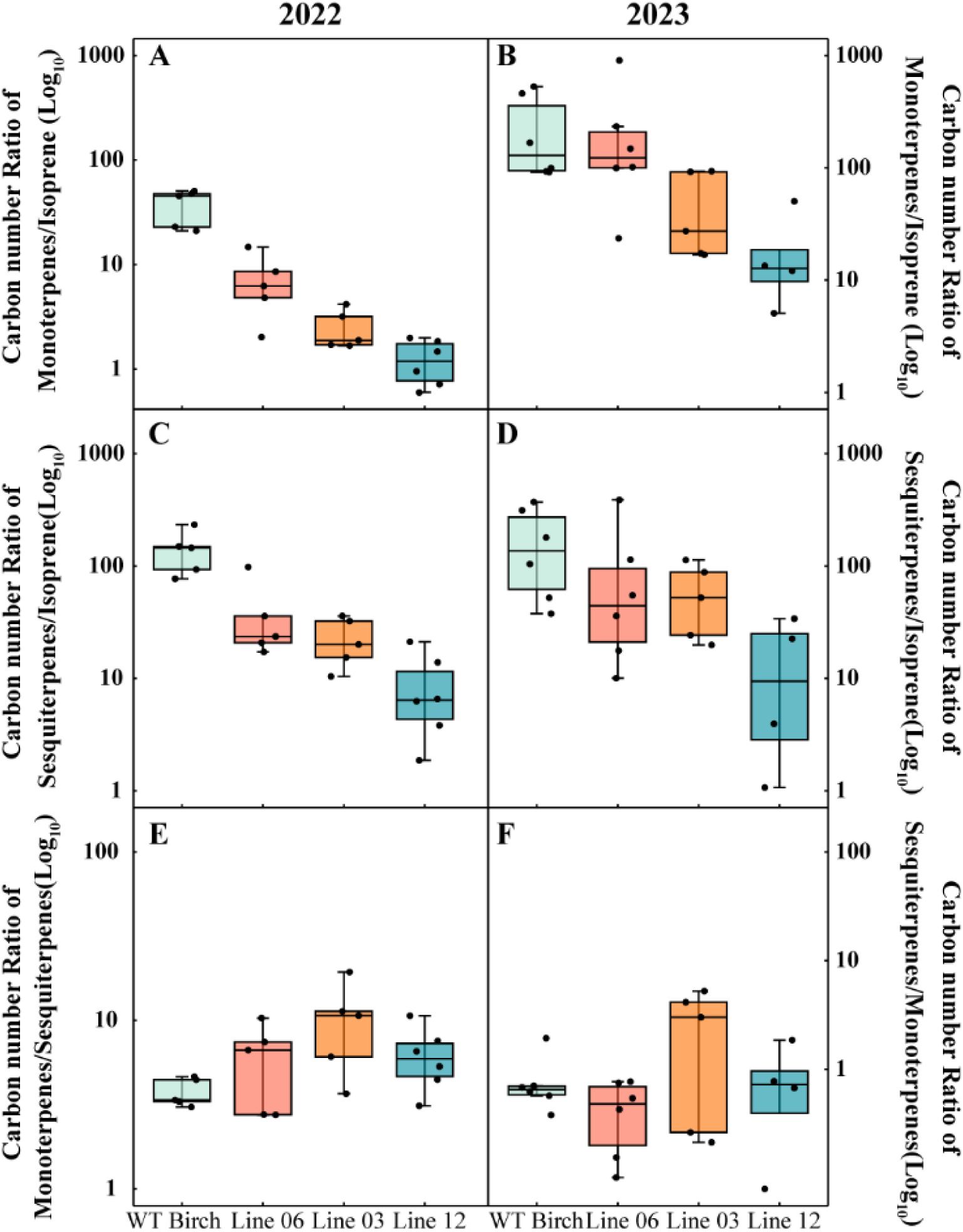
Carbon number ratios of terpenoid emissions from wild-type and transgenic silver birch lines in 2022 and 2023. Carbon number ratios of different terpenoid classes emitted by wild-type (WT) birch and three transgenic lines (06, 03, and 12) over two consecutive years. (A, B) Monoterpenes to isoprene ratio. (C, D) Sesquiterpenes to isoprene ratio. (E, F) Sesquiterpenes to monoterpenes ratio. Note the difference in scale. The boxes represent the interquartile range (IQR), with the median indicated by the horizontal line. Whiskers extend to 1.5 times the IQR, and data points beyond this range are shown as individual dots (*N* ≥ 4).

**Fig. S3.**
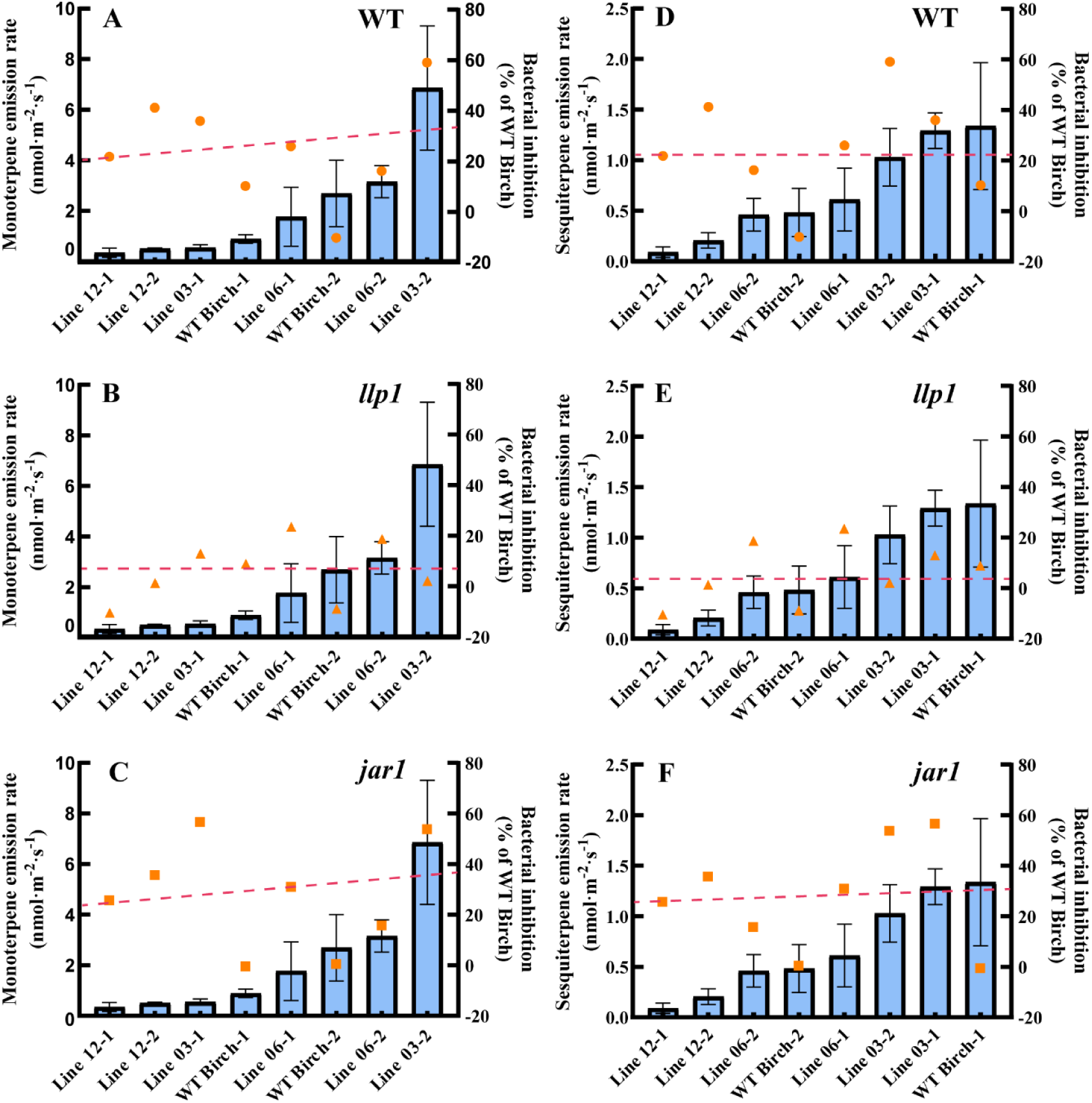
Monoterpene and sesquiterpene emission rates in birch lines and their correlation with bacterial growth inhibition in *Arabidopsis* mutants. Emission rates of (A-C) monoterpenes and (D-F) sesquiterpenes were measured in different birch lines triangle and tested against (A, D) wild-type (WT) Arabidopsis, (B, E) salicylic acid signaling-deficient llp1 mutant, and (C, F) jasmonic acid signaling-deficient jar1 mutant. Blue bars represent VOC emission rates from birch lines (mean ± SEM, *N* = 3). Orange symbols indicate bacterial growth inhibition in Arabidopsis genotypes relative to WT control (%). Red dashed lines illustrate the trend between VOC emissions and antimicrobial activity. Correlation analysis showed no significant relationships for monoterpenes [monoterpenes: WT (*R²* = 0.1212, *P* = 0.3980), *llp1* (*R²* = 0.009380, *P* = 0.8195), *jar1* (*R²* = 0.04996, *P* = 0.5947)] and sesquiterpenes [WT (*R²* = 0.02893, *P* = 0.6872), *llp1* (*R²* = 0.1343, *P* = 0.3719), *jar1* (*R²* = 0.03493, *P* = 0.6577)]. Numbers following birch line designations represent different emission triangles.

**Fig. S4.**
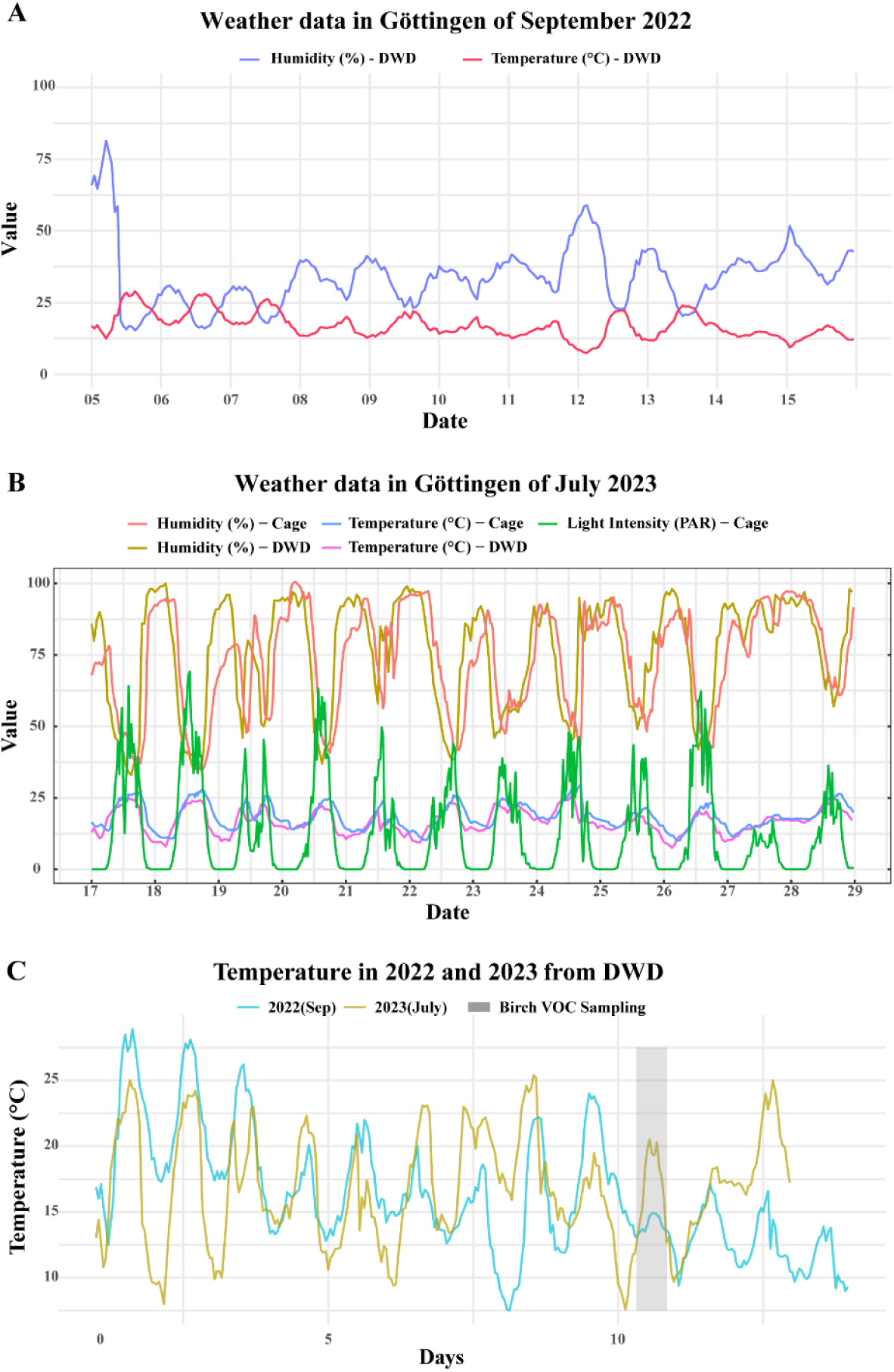
Environmental conditions during experimental periods in 2022 and 2023. (A) Temperature (°C, pink line) and relative humidity (%, yellow/green line) measured at the DWD (Deutscher Wetterdienst, German Weather Forcast Service) weather station in Göttingen from September 5–15, 2022. (B) Daily temperature (°C, blue lines), relative humidity (%, red/yellow lines), and light intensity (PAR, green line) recorded in the experimental cage and at the DWD weather station from July 17–29, 2023. (C) Comparison of temperature trends between September 2022 (light blue line) and July 2023 (orange line) from DWD data over a 12-day period. The grey shaded area marks the timing of birch VOC sampling. The temperature data illustrates temporal variation and differences in thermal conditions between the two experimental periods.

**Table S1.** List of Isoprene, monoterpenoids and sesquiterpenoids identified in birch lines with their retention times and Kovats index.

| Num | Compound | Retention Time | CAS Number | Kovats Index | Classification |
| --- | --- | --- | --- | --- | --- |
| 1 | Isoprene | 5.710 | 78-79-5 | 705 | Isoprene |
| 2 | α-Pinene | 9.668 | 80-56-8 | 894 | Monoterpenoids |
| 3 | D-limonene | 11.519 | 5989-27-5 | 982 | Monoterpenoids |
| 4 | β-Cyclocitral | 15.307 | 432-25-7 | 1164 | Monoterpenoids |
| 5 | (-)-Car-3-en-2-one | 16.120 | 53585-45-8 | 1202 | Sesquiterpenoids |
| 6 | Trans-α-Bergamotene | 19.538 | 13474-59-4 | 1366 | Sesquiterpenoids |
| 7 | 7-epi-Sesquithujene | 18.554 | 159407-35-9 | 1319 | Sesquiterpenoids |
| 8 | Gamma-Elemene | 19.654 | 29873-99-2 | 1371 | Sesquiterpenoids |
| 9 | Valerena-4,7(11)-diene | 19.694 | 351222-66-7 | 1373 | Sesquiterpenoids |
| 10 | Valencene | 19.978 | 4630-07-3 | 1387 | Sesquiterpenoids |
| 11 | α-Farnesene | 20.775 | 502-61-4 | 1425 | Sesquiterpenoids |
| 12 | β-Selinene | 20.900 | 17066-67-0 | 1431 | Sesquiterpenoids |
| 13 | SQT1 | 19.908 | - | 1383 | Sesquiterpenoids |
| 14 | SQT2 | 20.949 | - | 1433 | Sesquiterpenoids |
| 15 | SQT3 | 21.448 | - | 1457 | Sesquiterpenoids |
| 16 | Sabinene | 10.269 | 3387-41-5 | 923 | Monoterpenoids |
| 17 | β-Myrcene | 10.434 | 123-35-3 | 931 | Monoterpenoids |
| 18 | Trans-β-Ocimene | 11.249 | 3779-61-1 | 970 | Monoterpenoids |
| 19 | β-Ocimene | 11.488 | 13877-91-3 | 981 | Monoterpenoids |
| 20 | Linalool | 12.709 | 78-70-6 | 1039 | Monoterpenoids |
| 21 | (E)-4,8-Dimethylnona-1,3,7-triene | 12.994 | 19945-61-0 | 1053 | Monoterpenoids |
| 22 | Neo-allo-ocimene | 13.408 | 7216-56-0 | 1073 | Monoterpenoids |
| 23 | Allo-ocimene | 13.823 | 57396-75-5 | 1093 | Monoterpenoids |
| 24 | α-Cubebene | 17.889 | 17699-14-8 | 1287 | Sesquiterpenoids |
| 25 | Ylangene | 18.386 | 14912-44-8 | 1311 | Sesquiterpenoids |
| 26 | (-)-β-Bourbonene | 18.706 | 5208-58-2 | 1326 | Sesquiterpenoids |
| 27 | α-Santalene | 19.261 | 512-61-8 | 1353 | Sesquiterpenoids |
| 28 | β-Copaene | 19.374 | 18252-44-3 | 1358 | Sesquiterpenoids |
| 29 | α-Bergamotene | 19.496 | 18252-46-5 | 1364 | Sesquiterpenoids |
| 30 | Sesquisabinene | 19.622 | 58319-04-3 | 1370 | Sesquiterpenoids |
| 31 | Cis-β-Farnesene | 19.686 | 28973-97-9 | 1373 | Sesquiterpenoids |
| 32 | 6,9-Guaiadene | 19.769 | 36577-33-0 | 1377 | Sesquiterpenoids |
| 33 | β-Santalene | 19.845 | 511-59-1 | 1380 | Sesquiterpenoids |
| 34 | Sesquisabinene B | 20.032 | 1431636-59-7 | 1389 | Sesquiterpenoids |
| 35 | β-Guaiene | 20.185 | 88-84-6 | 1397 | Sesquiterpenoids |
| 36 | 1,3,6,10-Dodecatetraene,<br>3,7,11-trimethyl-, (Z,E)- | 20.419 | 26560-14-5 | 1408 | Sesquiterpenoids |
| 37 | β-Bergamotene | 20.599 | 15438-94-5 | 1416 | Sesquiterpenoids |
| 38 | Germacrene D | 20.683 | 23986-74-5 | 1420 | Sesquiterpenoids |
| 39 | β-Bisabolene | 20.982 | 495-61-4 | 1435 | Sesquiterpenoids |
| 40 | Sesquicineole | 21.174 | 90131-02-5 | 1444 | Sesquiterpenoids |
| 41 | γ-Cadinene | 21.307 | 39029-41-9 | 1450 | Sesquiterpenoids |
| 42 | β-sesquiphellandrene | 21.365 | 20307-83-9 | 1453 | Sesquiterpenoids |
| 43 | Sesquirosefuran | 21.725 | 39007-93-7 | 1470 | Sesquiterpenoids |
| 44 | 7-epi-trans-Sesquisabinene<br>hydrate | 21.841 | 145512-84-1 | 1476 | Sesquiterpenoids |
| 45 | 7-epi-cis-Sesquisabinene<br>hydrate | 21.940 | 11028-42-5 | 1481 | Sesquiterpenoids |

